# Skeletal Muscle Metrics on Clinical ^18^F- FDG PET/CT Predict Health Outcomes in Patients with Sarcoma

**DOI:** 10.1101/304279

**Authors:** Brent Foster, Robert D. Boutin, Leon Lenchik, David Gedeon, Yu Liu, Vinay Nittur, Ramsey D. Badawi, Chin-Shang Li, Robert J. Canter, Abhijit J. Chaudhari

## Abstract

The aim of this study was to determine the association of measures of skeletal muscle determined from ^18^F-FDG PET/CT with health outcomes in patients with soft-tissue sarcoma. 14 patients (8 women and 6 men; mean age 66.5 years) with sarcoma had PET/CT examinations. On CTs of the abdomen and pelvis, skeletal muscle was segmented, and cross-sectional muscle area, muscle volume, and muscle attenuation were determined. Within the segmented muscle, intramuscular fat area, volume, and density were derived. On PET images the standardized uptake value (SUV) of muscle was determined. Regression analyses were conducted to determine the association between the imaging measures and health outcomes including overall survival (OS), local recurrence-free survival (LRFS), distant cancer recurrence (DCR), and major surgical complications (MSC). The association between imaging metrics and pre-therapy levels of serum C-reactive protein (CRP), creatinine, hemoglobin, and albumin was determined. Decreased volumetric muscle CT attenuation was associated with increased DCR. Increased PET SUV of muscle was associated with decreased OS and LRFS. Lower muscle SUV was associated with lower serum hemoglobin and albumin. Muscle measurements obtained on routine ^18^F-FDG PET/CT is associated with outcomes and serum hemoglobin and albumin in patients with sarcoma.

## INTRODUCTION

Skeletal muscle has a four times higher storage capacity of glycogen compared to the liver and plays a critical role in glycemic control.[1] In metabolic homeostasis, skeletal muscle accounts for 30% of the resting metabolic rate and 80% of glucose disposal under insulin stimulated conditions.[2]

Cachexia (wasting of muscle) occurs in more than half of all cancer patients and is strongly correlated with adverse patient outcomes, including death.[3–5] Poor nutritional intake, competition with metabolically demanding tumor(s), or alterations in resting metabolism cannot fully explain the loss of muscle mass seen in patients with cancer. The pathogenesis of cachexia in patients with cancer may include the production of acute phase response proteins that require catabolism of muscle. [6] Although acute phase response proteins such as C-reactive protein (CRP) and albumin are commonly obtained in routine blood testing [7], their relationship with muscle wasting is unclear.

^18^F-FDG PET/CT scans are routinely acquired for staging of cancer patients. Important muscle metrics may be derived opportunistically from these scans to help determine prognosis. CT-measured muscle metrics have been used for prognosis in various types of cancer patients. In particular, fatty infiltration of muscle, i.e., increased intramuscular adipose tissue (IMAT), has been associated with increased morbidity and mortality.[8, 9] PET-based muscle metrics have the potential for adding complementary information to CT measures for prognosis but have not yet been investigated.

The purpose of our study was to (1) evaluate the association of CT and PET muscle metrics with overall survival (OS), local recurrence-free survival (LRFS), distant cancer recurrence (DCR), and major surgical complications (MSC) in patients with sarcoma and (2) determine the associations of the imaging metrics with serum CRP, creatinine, hemoglobin, and albumin.

## METHODS

### Patient population

Approval for this study was obtained from the Institutional Review Board, and the requirement for informed consent was waived due to the retrospective nature of the study. Participants in this study were part of a larger cohort with biopsy proven soft-tissue sarcoma who received treatment between January 2008 and February 2013. Inclusion criteria were (1) a whole-body ^18^F-FDG PET/CT obtained for disease staging prior to starting treatment and (2) blood tests obtained within one month of the ^18^F-FDG PET/CT scan prior to treatment. We only included patients imaged at our institution to reduce heterogeneity in image acquisition parameters and documentation of health related outcomes. Sixteen patients met our inclusion criteria. Two patients were excluded due to misregistration between PET and CT images due to motion. In the remaining 14 patients (8 female and 6 male) PET/CT scans were analyzed. The clinical outcome variables were obtained from the electronic medical records: OS (in months), LRFS (in months), DCR (yes/no), and MSC (yes/no) as well as serum biomarkers CRP (mg/L), creatinine (mg/dL), hemoglobin (g/dL), and albumin (g/dL). Table 1 gives a description of the patient characteristics, serum biomarkers, and outcome measures.

**TABLE 1.**
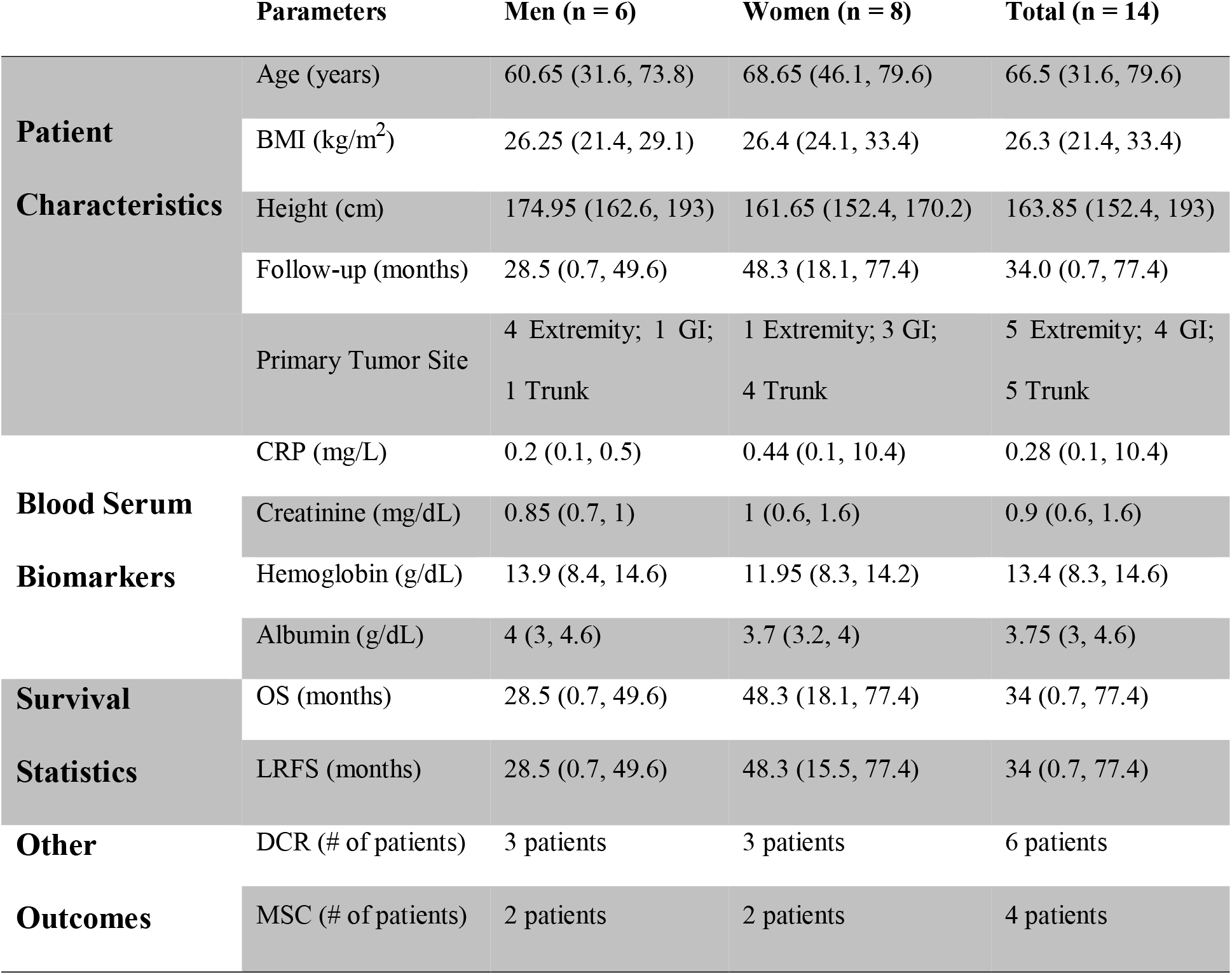
Patient Characteristics, Serum Biomarkers, and Outcome Measures

Data presented is the median with the range in parentheses.

**Abbreviations:** BMI, Body mass index; GI, Gastrointestinal; CRP, C-reactive protein; OS, Overall survival; LRFS, Local recurrence-free survival; DCR, Distant cancer recurrence; MSC, Major surgical complications.

### Image acquisition

The PET/CT images were acquired on a PET/CT whole body scanner (Discovery ST, GE Healthcare). The CT was acquired in helical mode with a voxel size of 0.98×0.98×3.7 mm, and with a 140 kVp for 12 patients and 120 kVp for 2 patients. For PET, data were acquired in 2D mode. An injected dose of 761.09±100.27 MBq (i.e. 20.6±2.7 mCi) was used and the time from injection to imaging was 71.77±14.78 minutes. Reconstructed PET images (voxel size of 5.47×5.47×3.27 mm) incorporating all image corrections used the manufacturer’s software and recommended reconstruction parameters(ordered-subset expectation maximization with 30 subsets, 2 iterations, and post-reconstruction filter with 7.0 mm full width at half maximum).

### CT segmentation

In order to standardize the volume of muscle segmented, CT scans were cropped from the level of vertebral pedicle at T12 to the ischial tuberosity (Fig. 1). After cropping, all muscle tissue was segmented using CT attenuation thresholds: −29 to 150 HU for muscle and −190 to −30 HU for intramuscular fat.[10] Manual segmentation was used to correct for any errors seen after thresholding (Fig. 2). The cropping was done using 3D Slicer (www.slicer.org) while the manual segmentation was done using the Brainsuite software (www.brainsuite.org).

**FIGURE 1:**
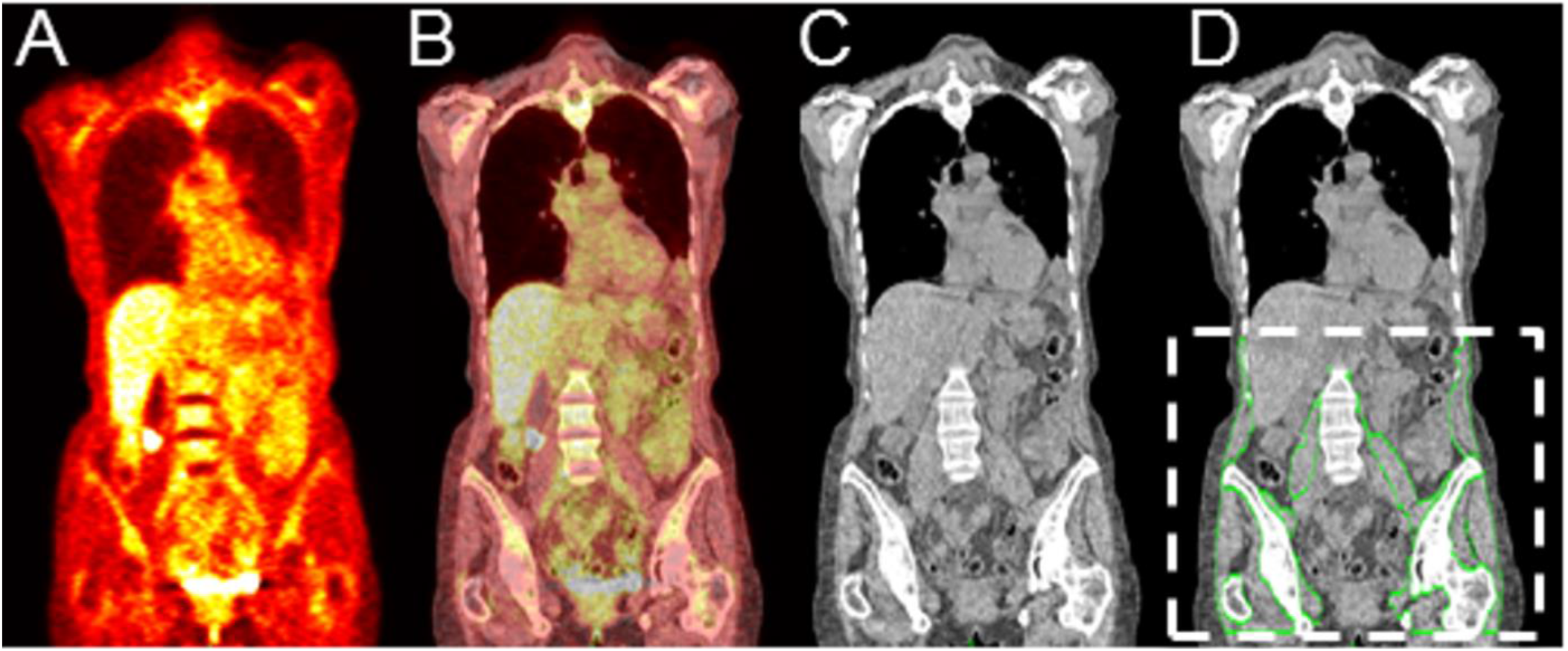
(A) PET, (B) fused PET/CT, and (C) CT coronal section of a representative patient. The segmented skeletal muscle is outlined by green lines and overlaid onto the corresponding CT coronal section (D). The dotted white lines in (D) show the standardized body region used for segmentation in our study.

**FIGURE 2:**
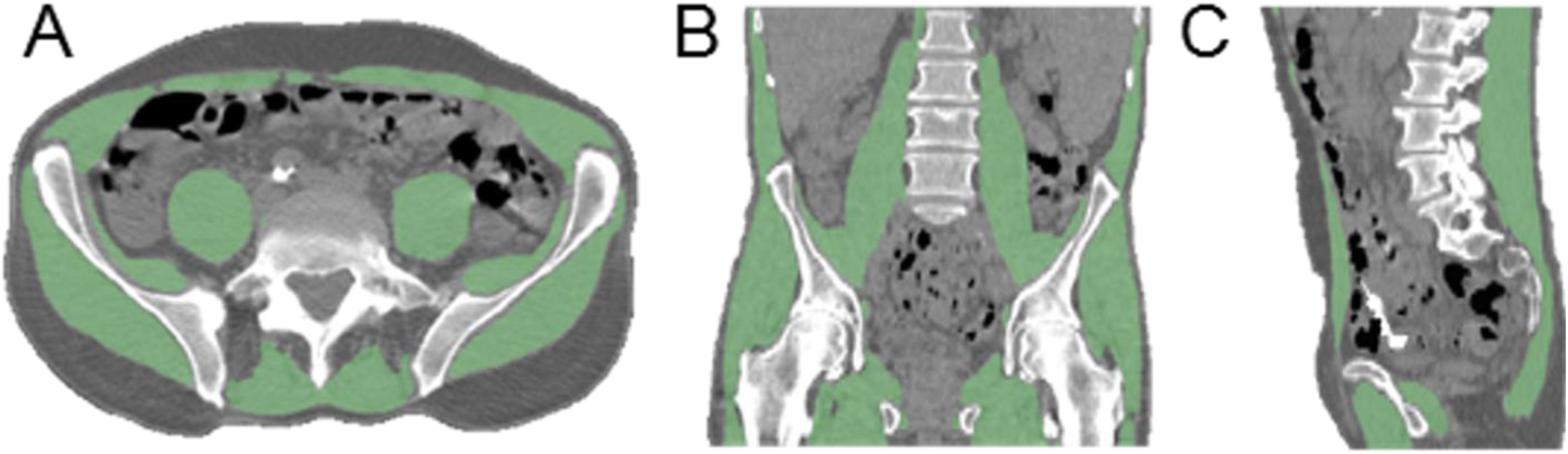
Skeletal muscle segmentation; (A) Axial, (B) Coronal, and (C) Sagittal sections from CT (gray scale) with overlaid segmented skeletal muscle (green areas).

The axial section location was identified from the sagittal view of the spine on CT, and the centers of the L3 and L4 vertebrae were identified. Using the muscle mask that was previously created volumetrically, axial cross-sections at the L3 and L4 levels were extracted.

Imaging metrics for both the cross-sectional (2D) and volumetric (3D) analysis of muscle were computed using MATLAB (2014 64-bit version) on a Windows 7 PC with 32.0 GB of RAM and a 3.50 GHz processor. Table 2 gives a description of the imaging metrics that were calculated in analyzing muscle quantity and quality. For the two cross-sections, 2D equivalents of the 3D imaging measures were derived (i.e., area in cm^2^ instead of volume in cm^3^).

**TABLE 2.** Imaging Measures Derived

| Measure type | CT |  |  | PET |  |  |
| --- | --- | --- | --- | --- | --- | --- |
|  | Measures | Men | Women | Measures | Men | Women |
| **Volume Measures (Liters)** | Body | 20.8 (18.4, 33.6) | 22.6 (16.0, 30.4) |  | N/A | N/A |
|  | Muscle | 7.5 (6.2, 9.6) | 4.8 (4.2, 6.2) | Muscle MAV | 0.2 (0, 0.8) | 0.2 (0, 1.6) |
|  | IMAT | 1.9 (0.9, 2.6) | 1.4 (0.96, 3.1) | IMAT MAV | 0.1 (0, 0.3) | 0.1 (0, 0.6) |
| **Muscle Measures (HU or SUB)** | Max | 1026 (491, 3071) | 981 (363, 1124) | Max | 4.7 (3.1, 5.8) | 4.5 (3.4, 5.5) |
|  | Mean | 29.3 (14.4, 49.4) | 21.9 (4.8, 37.3) | Mean | 0.8 (0.7, 1.2) | 1.0 (0.9, 1.2) |
|  | SD | 20.3 (14.9, 24.1) | 16.4 (11.2, 23.7) | SD | 0.2 (0.1, 0.4) | 0.3 (0.2, 0.3) |
| **IMAT Measures (HU or SUB)** | Mean | -30.3 (-52.5, -14.3) | -31.3 (-53.5, -13.6) | Mean | 0.8 (0.6, 1.2) | 0.9 (0.8, 1.1) |
|  | SD | 39.3 (18.0, 56.1) | 34.8 (16.3, 61.3) | SD | 0.4 (0.2, 0.5) | 0.4 (0.2, 0.5) |

Data presented is the median with the range in parentheses, and PET measures are using the SUB standardization.

**Abbreviations:** MAV, Metabolically active volume; IMAT, Intramuscular adipose tissue; HU, Hounsfield Units; SUB, Standardized uptake value normalized by body weight.

### PET image processing

The PET images were interpolated to the CT image resolution using standard bicubic interpolation. The PET images were then cropped using the same bounding box as the CT images, and the masks for muscle and IMAT were applied. The PET images were standardized using the two most common standardized uptake value (SUV) definitions based on the total body weight (SUB) and based on an estimation of the lean body mass utilizing the patient’s sex and height (SUL). [11] The SUV quantification was validated by analyzing the mean and standard deviation of each definition within a manually defined region of interest in the liver and comparing with accepted values in the literature.[11] For the study population, the SUBmean was found to be 2.67±0.48 while the SULmean was 2.25±0.37. Potential confounding in the SUV measures caused by different radiotracer uptake times in patients was evaluated. No relationship was found between the SUBmean or SULmean of the liver versus the radiotracer uptake time (p=0.61). Due to the intensity normalization step and the relatively slow washout of ^18^F-FDG in muscle, the variation in uptake time was not expected to significantly change the SUB or SUL.[12]

### Statistical analysis

Cox proportional hazards regression[13] was used to determine the association of imaging metrics with time-to-event outcome variables, while logistic regression was employed when the outcome variables were binary. Robust linear regression was used to determine the association between imaging metrics and the serum biomarkers. The patient’s age at diagnosis was controlled for in al l statistical analyses. A p-value of <0.05 was considered statistically significant. A marginally significant relationship was defined as 0.05<p<0.10. Analyses were performed using SAS v9.4 (SAS Institute Inc., Cary, NC, USA).

To evaluate the agreement between cross-sectional volumetric muscle measures the mean absolute percent difference in the values was computed. The agreement between the cross-sectional measures at the L3 and L4 levels was also calculated.

## RESULTS

### Cross-sectional versus volumetric measures

The agreement of the volumetric measures with the cross-sectional measures is provided in Table 3 with the values given in mean ± SD. The SD in this case indicates the variation of the mean across subjects. The most well conserved measure was the SUB_mean_ and the SUL_mean_ with about a 13% difference between the two cross-sections and the volumetric measures, and with less than 4% difference between the L3 and L4 cross-sections. The least conserved variable was measured on CT: the HU_max_ (>127 to 139 HU when comparing the volumetric and cross-sections). Furthermore, there was a large variation of the percent IMAT in muscle observed with CT between the volumetric and crosssections (62 to 65%).

**TABLE 3.** Agreement Between Cross-sectional and Volumetric Imaging Measures

| Imaging measures | L3 versus L4 | L3 versus Volumetric | L4 versus Volumetric |
| --- | --- | --- | --- |
| **HU<sub>max</sub>** | 26.4 ± 26.4 | 139 ± 27.8 | 127.3 ± 26 |
| **HU<sub>mean</sub>** | 22.9 ± 27.5 | 29.6 ± 25.3 | 39.4 ± 45.1 |
| **HU<sub>SD</sub>** | 10.3 ± 16.1 | 65.3 ± 21.4 | 67.1 ± 24.9 |
| **Percent Muscle of Body** | 6.2 ± 6.4 | 15.8 ± 8.6 | 15.8 ± 8.2 |
| **IMAT Percent of Muscle** | 5.9 ± 2.9 | 64.9 ± 40.2 | 61.6 ± 41.9 |
| **SUB<sub>max</sub>** | 17.3 ± 12.5 | 52.5 ± 21.7 | 46 ± 17 |
| **SUB<sub>mean</sub>** | 3.4 ± 2.4 | 13.2 ± 8.7 | 12.6 ± 8.6 |
| **SUB<sub>SD</sub>** | 8.6 ± 6.5 | 23.2 ± 18.9 | 27.5 ± 17.5 |
| **SUL<sub>max</sub>** | 15.9 ± 10.7 | 28.2 ± 16.7 | 33 ± 11.6 |
| **SUL<sub>mean</sub>** | 3.4 ± 2.4 | 12.3 ± 7.8 | 11.6 ± 7.9 |
| **SUL<sub>SD</sub>** | 5.7 ± 4.4 | 18.3 ± 11.3 | 19.8 ± 13.5 |

Measures are the average absolute percent difference in mean ± standard deviation (SD) across the subjects.

**Abbreviations:** L3, 3rd lumbar vertebra level; L4, 4th lumbar vertebra level; HU, Hounsfield Units; IMAT, Intramuscular adipose tissue; SUB, Standardized uptake value normalized by body weight; SUL, Standardized uptake value normalized by lean body mass.

### Muscle attenuation and metabolic volume/area

The HU_mean_ from CT, volumetrically and for the individual cross-sections, had a negative linear relationship with the metabolically active volume/area for muscle. The IMAT volume/area from CT did not show an association with the PET measures.

### Cross-sectional muscle measures and outcomes

The SUL_mean_ of the IMAT at the L3 cross-section was significantly associated with the OS (hazard ratio (HR)=59.21, 95% CI=(1.29, 2720.91)) and LRFS (HR=39.83, 95% CI=(1.16, 1371.16)), after adjustment for age at diagnosis. More specifically, a higher SUL_mean_ of the IMAT at the L3 crosssection had a significantly shorter OS and a shorter LRFS. A similar trend was observed for the SUL_mean_ of the IMAT at the L4 level; this measure was marginally significantly associated with the OS (HR=82.38, 95% CI=(0.96, 7104.32), p=0.052) and LRFS (HR=55.66, 95% CI=(0.93, 3345.49), p=0.055). The cross-sectional measures based on CT were not associated with patient outcomes.

### Volumetric muscle measures and outcomes

Higher volumetric HU_mean_ of muscle from CT was marginally significantly associated with a shorter LRFS (HR=1.06, 95% confidence interval (CI)=(0.99, 1.13), p=0.088). In addition, higher volumetric HU standard deviation (HUSD) was marginally significantly associated with lower odds of DCR (odds ratio (OR)=0.90, 95% CI=(0.79, 1.02), p=0.085).

A significant difference between the HU_max_ of the DCR group and the no DCR group implied that patients with lower HU_max_ were more likely to have DCR while those with a higher HU_max_ tended to have a lower rate of DCR. OS and MSC did not associate with any CT volumetric measure. None of the volumetric measures from PET were associated with the outcomes studied.

### Muscle measures and serum biomarkers

#### Creatinine

Measures that had a positive relationship with creatinine level were the volumetric IMAT metabolically active volume, and the HU_sd_, both volumetrically-derived and from the two crosssections. Volumetric and cross-sectional HU_mean_ had an inverse relationship with creatinine. None of the cross-sectional measures from PET were associated with creatinine.

### CRP

The body volume from CT, the percent IMAT in the muscle volume, the SUB_sd_ of the IMAT at the L3 level and muscle HU_sd_ at the L4 level had a positive linear relationship with the CRP level. The HU_max_ at L3 and HUmean at L4 had an inverse relationship with CRP. None of the volumetric imaging measures from PET were associated with CRP.

### Hemoglobin

Both volumetric and cross-sectional measures from HU showed no association with the hemoglobin level. SUB_mean_ of muscle and SUB_mean_ of the IMAT, measured both volumetrically and for both cross-sections, had a negative linear relationship with hemoglobin. Additionally, the SUB_max_, SUL_max_, SUB_sd_, and SUL_sd_ of muscle for the L3 cross-section also had a negative linear relationship with hemoglobin.

### Albumin

A range of PET measures, both volumetric and cross-sectional, showed a negative linear relationship with albumin. These include SUB_mean_, SUB_sd_, SUL_sd_, SUB_mean_ of the IMAT, and SUBsd of the IMAT. Additionally, the metabolically active area of muscle and IMAT at the L3 and L4 levels showed a negative linear relationship with albumin. The volumetric and cross-sectional measures from CT showed no association with albumin.

## DISCUSSION

^18^F-FDG PET/CT images are routinely acquired for cancer staging, with the primary focus being on tumor assessment and evaluation of the extent of malignant spread. The scan can also be used to opportunistically measure muscle metrics, important for the evaluation of cancer-related cachexia.

Our findings in sarcoma patients show that ^18^F-FDG PET/CT measures are associated with important health outcomes and serum biomarkers. These imaging biomarkers may complement clinical examination of muscle and aid in therapy selection and evaluation. The opportunistic biomarkers provide an alternative to other techniques used for assessing cachexia such as dual x-ray absorptiometry (DXA), bioimpedance analysis (BIA), and functional testing (e.g., handgrip strength and gait speed).[14] Ultimately, these PET/CT biomarkers could help provide insight into the mechanisms of cancer cachexia, and help in identifying disease targets or therapies.

Prior studies of muscle depletion have typically used single slice CT images at L3 or L4, but we are not aware of comparisons between these two levels or with volumetric analysis.[15–18] Martin et al.[16] used two axial sections from CT, both taken at the L3 vertebral level while Taguchi et al.[17] used a single axial cross-section at the L3 level. Tsein et al.[17] used a single axial section at the middle of the L4 level. Baumgartner et al.[15] assessed six abdominal cross-sections of clinically normal patients, with the top section located at the caudal tip of the xiphoid process while the bottom section located at the cranial edge of the iliac crests. ^18^F-FDG PET-based measures of muscle have not been evaluated for predicting outcomes in patients with cancer.

Our study derived volumetric measures from both CT and PET, thus surveying large parts of the body. We believe that volumetric measures will be more reproducible and robust compared to crosssectional area measures (in our case, from the L3 and L4 cross-sections), analogous to findings regarding tumor volumetric measurements.[19] Our results show that imaging measures from a single axial section may not be representative of measures derived using volumetric analyses. Overall, more volumetric measures were associated with health outcomes compared with cross-sectional measures.

A higher level of IMAT has been associated with lower muscle quality resulting in muscle fatigue, bone fragility, and disability.[8, 9] IMAT in combination with muscle depletion, characterized by a reduction in muscle size, has been measured using CT.[15–18] Our study found that a higher HUmean correlated with lower LRFS. Muscle with a higher HUmean would have less IMAT. This finding suggests that patients with more IMAT have better outcomes, possibly due to the “obesity paradox”.[20] The obesity paradox is the observation that although obese patients are more likely to get cancer they also tend to have better survival. In separate analysis (not shown), BMI in our cohort was not a predictor of overall survival (p=0.25) or other survival outcomes.

We examined the relationships between imaging measures and four commonly used serum biomarkers. As in cachexia, patients with age-related muscle wasting have high serum CRP[21], low hemoglobin levels[21], and low serum albumin[22]. Frail patients also have high serum CRP[23]. A positive relationship has been found between creatinine and lean muscle mass as measured by DXA.[24] In our study, increased IMAT was associated with high creatinine and CRP levels. Our results show that higher metabolic measures of muscle from PET were associated with lower serum albumin and lower hemoglobin levels. The predictive power of serum biomarkers for cancer outcomes and patient frailty are actively debated[25], and imaging associations found in this paper may help clarify their implications.

There are several limitations of our study. Our sample size was small, and the associations measured must be tested in a larger cohort of patients. Although we used the two most common definitions of SUV, normalized by body weight and lean body mass, it has also been proposed to use glucose corrected SUV measures.[26] Lastly, manual volumetric muscle segmentation is fairly time consuming. A potential solution to this problem is to combine our existing muscle segmentation masks to construct a statistical muscle atlas. This atlas could then be warped to additional patient CT images using deformable registration to automatically segment muscle with machine learning in a computationally efficient reproducible manner. Such methodological advancements would allow for processing images of a larger patient cohort while still retaining the advantages of volumetric analysis.

## CONCLUSION

Metabolic information from ^18^F-FDG PET may complement that gained from CT for the characterization of muscle. Several muscle biomarkers could be used as predictors of health outcomes.

## ACKNOWLEDGMENTS

This project was supported by the National Institutes of Health grants R03EB015099, K12HD051958, and the National Center for Advancing Translational Sciences (NCATS) grant #UL1 TR000002. The content is solely the responsibility of the authors and does not necessarily represent the official views of the National Institutes of Health. The funders had no role in study design, data collection and analysis, decision to publish, or preparation of the manuscript.

## REFERENCES

1. Berg, J.M., J.L. Tymoczko, and L. Stryer, in Biochemistry. 2002, W H Freeman.

2. Egan, B. and J.R. Zierath, Exercise metabolism and the molecular regulation of skeletal muscle adaptation. Cell Metab, 2013. 17(2): p. 162-84.

3. Acharyya, S., et al., Dystrophin glycoprotein complex dysfunction: a regulatory link between muscular dystrophy and cancer cachexia. Cancer Cell, 2005. 8(5): p. 421-32.

4. Evans, W.J., et al., Cachexia: a new definition. Clin Nutr, 2008. 27(6): p. 793-9.

5. Fearon, K., et al., Definition and classification of cancer cachexia: an international consensus. Lancet Oncol, 2011. 12(5): p. 489-95.

6. Bonetto, A., et al., STAT3 activation in skeletal muscle links muscle wasting and the acute phase response in cancer cachexia. PLoS One, 2011. 6(7): p. e22538.

7. Cray, C., J. Zaias, and N.H. Altman, Acute phase response in animals: a review. Comp Med, 2009. 59(6): p. 517-26.

8. Malafarina, V., et al., Sarcopenia in the elderly: diagnosis, physiopathology and treatment. Maturitas, 2012. 71(2): p. 109-14.

9. Manini, T.M., et al., Reduced physical activity increases intermuscular adipose tissue in healthy young adults. Am J Clin Nutr, 2007. 85(2): p. 377-84.

10. Mourtzakis, M., et al., A practical and precise approach to quantification of body composition in cancer patients using computed tomography images acquired during routine care. Appl Physiol Nutr Metab, 2008. 33(5): p. 997-1006.

11. Paquet, N., et al., Within-patient variability of 18F-FDG: standardized uptake values in normal tissues. Journal of Nuclear Medicine, 2004. 45(5): p. 784-788.

12. Goodpaster, B.H., et al., Interactions Among Glucose Delivery, Transport, and Phosphorylation That Underlie Skeletal Muscle Insulin Resistance in Obesity and Type 2 Diabetes: Studies With Dynamic PET Imaging. Diabetes, 2014. 63(3): p. 1058-1068.

13. Cox D.R., Regression Models and Life-Tables. Journal of the Royal Statistical Society. Series B (Methodological), 1972. 34(2): p. 187-220.

14. Cesari, M., et al., Biomarkers of sarcopenia in clinical trials-recommendations from the International Working Group on Sarcopenia. J Cachexia Sarcopenia Muscle, 2012. 3(3): p. 181-190.

15. Baumgartner, R.N., et al., Abdominal composition quantified by computed tomography. Am J Clin Nutr, 1988. 48(4): p. 936-45.

16. Martin, L., et al., Cancer cachexia in the age of obesity: skeletal muscle depletion is a powerful prognostic factor, independent of body mass index. J Clin Oncol, 2013. 31(12): p. 1539-47.

17. Tsien, C., et al., Reversal of sarcopenia predicts survival after a transjugular intrahepatic portosystemic stent. Eur J Gastroenterol Hepatol, 2013. 25(1): p. 85-93.

18. Taguchi, S., et al., Sarcopenia evaluated by skeletal muscle index is a significant prognostic factor for metastatic urothelial carcinoma. Clinical Genitourinary Cancer, 2015.

19. Frenette, A., et al., Do Diametric Measurements Provide Sufficient and Reliable Tumor Assessment? An Evaluation of Diametric, Areametric, and Volumetric Variability of Lung Lesion Measurements on Computerized Tomography Scans. Journal of Oncology, 2015. 2015.

20. Amundson, D.E., S. Djurkovic, and G.N. Matwiyoff, The obesity paradox. Crit Care Clin, 2010. 26(4): p. 583-96.

21. Ida, S., et al., Sarcopenia is a Predictor of Postoperative Respiratory Complications in Patients with Esophageal Cancer. Annals of surgical oncology, 2015: p. 1-6.

22. Baumgartner, R.N., et al., Serum albumin is associated with skeletal muscle in elderly men and women. American Journal of Clinical Nutrition, 1996. 64(4): p. 552-558.

23. Cesari, M., et al., Frailty syndrome and skeletal muscle: results from the Invecchiare in Chianti study. Am J Clin Nutr, 2006. 83(5): p. 1142-8.

24. Nyasavajjala, S.M., et al., Creatinine and myoglobin are poor predictors of anaerobic threshold in colorectal cancer and health. Journal of Cachexia, Sarcopenia and Muscle, 2015. 6(2): p. 125-131.

25. Sawyers C.L., The cancer biomarker problem. Nature, 2008. 452(7187): p. 548-52.

26. Nozawa, A., et al., Glucose corrected standardized uptake value (SUVgluc) in the evaluation of brain lesions with 18F-FDG PET. European journal of nuclear medicine and molecular imaging, 2013. 40(7): p. 997-1004.

